# The impact of non-neutral synonymous mutations when inferring selection on non-synonymous mutations

**DOI:** 10.1101/2024.02.07.579314

**Authors:** Aina Martinez i Zurita, Christopher C. Kyriazis, Kirk E. Lohmueller

**Affiliations:** Department of Human Genetics, David Geffen School of Medicine, University of California, Los Angeles, USA; Interdepartmental Program in Bioinformatics, University of California, Los Angeles, USA; Department of Ecology and Evolutionary Biology, University of California, Los Angeles, USA

## Abstract

The distribution of fitness effects (DFE) describes the proportions of new mutations that have different effects on reproductive fitness. Accurate measurements of the DFE are important because the DFE is a fundamental parameter in evolutionary genetics and has implications for our understanding of other phenomena like complex disease or inbreeding depression. Current computational methods to infer the DFE for nonsynonymous mutations from natural variation first estimate demographic parameters from synonymous variants to control for the effects of demography and background selection. Then, conditional on these parameters, the DFE is then inferred for nonsynonymous mutations. This approach relies on the assumption that synonymous variants are neutrally evolving. However, some evidence points toward synonymous mutations having measurable effects on fitness. To test whether selection on synonymous mutations affects inference of the DFE of nonsynonymous mutations, we simulated several possible models of selection on synonymous mutations using SLiM and attempted to recover the DFE of nonsynonymous mutations using Fit∂a∂i, a common method for DFE inference. Our results show that the presence of selection on synonymous variants leads to incorrect inferences of recent population growth. Furthermore, under certain parameter combinations, inferences of the DFE can have an inflated proportion of highly deleterious nonsynonymous mutations. However, this bias can be eliminated if the correct demographic parameters are used for DFE inference instead of the biased ones inferred from synonymous variants. Our work demonstrates how unmodeled selection on synonymous mutations may affect downstream inferences of the DFE.

## Introduction

Mutations can have different effects on the fitness of an individual, ranging from a fitness benefit, to being neutral with respect to the chances of survival of the individual, or causing harmful or even lethal effects on the carrier. The distribution of fitness effects (DFE) is a probability distribution that describes the relative abundances of new mutations in each category (Eyre-Walker and Keightley, 2007). That is, the DFE describes the expected abundance of mutations with a particular fitness effect.

Understanding the DFE of new mutations is fundamental for both practical and theoretical questions. From a basic biology standpoint, the DFE directly describes how selection might be affecting the maintenance of genetic variation in a population. In addition, the DFE of a given species impacts our understanding of the molecular clock (Ohta, 1992) and has implications for the evolution of sex and recombination (Keightly and Otto, 2006). From a practical standpoint, having a reasonable estimate of the DFE for new mutations helps in understanding the amount of phenotypic variance a given mutation could explain, in turn affecting the genetic architecture of complex disease (Eyre-Walker, 2010; Eyre-Walker, 2006). In addition, DFE estimates can inform our decision making in the management of small populations of plants or animals in conservation efforts (Robinson et al. 2023; Kyriazis et al. 2021; Kyriazis et al. 2023). In summary, accurate DFE estimates for new mutations impact many fundamental aspects of modern population genetics.

There have been two main approaches used to estimate the DFE of new mutations. The first group of approaches uses direct experimental measurements applied to microorganisms, conducted either via mutagenesis experiments or mutation accumulation experiments (Bataillon and Bailey, 2014; Elena et al. 1998). Experimental approaches allow for direct measurement of the DFE. However, they are limited to organisms that are amenable to experimental manipulation, which severely limits the range of possible life forms for which the DFE can be estimated. Given the existing evidence of differences in the DFE between species (Huber et al. 2017), estimates from microorganisms are insufficient to properly characterize the DFE of higher-order species.

A second class of methods consists of computational approaches applied on polymorphism data derived from natural populations (Kim et al. 2017; Boyko et al 2008; Keightley and Eyre-Walker, 2007; Eyre-Walker et al. 2006; Tataru and Batallion, 2019). This suite of methods estimates the DFE that predicts the polymorphism data summarized by the site-frequency spectrum (SFS), or numbers of variants at different allele frequencies in the sample of individuals (Eyre-Walker et al. 2006, Kim et al. 2017, Keightley and Eyre-Walker, 2007, Tataru and Batallion, 2019) (Supplementary Figure 1). Using these methods, researchers have estimated the DFE of non-synonymous mutations in humans (Kim et al. 2017; Li et al. 2010; Boyko et al. 2008; Huang et al. 2021) as well as numerous other taxa including animals and plants (Castellano et al. 2019; Galtier 2016; Chen et al. 2017).

The SFS of non-synonymous variants is shaped by selective pressures as well as demographic processes (Williamson et al. 2005). Hence, methods need to control for demographic history in order to infer the DFE parameters. The majority of methods follow a similar workflow (Supplementary Figure 2). First, the genetic variation data is divided in two classes of sites: one putatively neutral and the other potentially experiencing the effects of selection (e.g. non-synonymous sites). Both categories are summarized into their respective SFS. Then, the neutral SFS is used to infer the effect of demographic processes that could cause changes in the SFS at all sites in the genome. Finally, conditioning the demographic history, a DFE is fitted to the SFS from the variants potentially experiencing selection. The deviation in the SFS of the sites putatively under selection from the SFS expected under the demographic model is used to fit the parameters of the DFE.

Historically, synonymous variants have been assumed to be effectively neutral since they lack an effect on the amino acid sequence (Williamson et al. 2005; Bailey et al. 2021). Following this logic, it is standard to use synonymous variation as a neutral reference to estimate demographic history prior to performing DFE inference (Boyko et al. 2008; Kim et al. 2017; Galtier 2016). The use of synonymous variants also helps account for background selection in between non-synonymous sites that could be skewing the non-synonymous SFS, since synonymous sites are interdigitated in the genome and therefore under the same background selection pressure (Kim et al. 2017).

However, several lines of evidence point towards synonymous mutations potentially experiencing selection. Experimental studies conducted in a range of microorganisms have found evidence of synonymous mutations causing both deleterious and beneficial selective effects (Bailey et al. 2021). Widespread evidence of codon bias across different taxa suggests synonymous mutations being impacted by selective forces (Hershberg and Petrov, 2008; Carlini and Stephan, 2003). Some synonymous mutations have been identified as having significant contributions to human disease (Sauna and Kimchi-Sarfaty, 2011), however, the overall impact of synonymous mutations on human disease may be minor (Dhindsa et al. 2022). Studies on genetic data from both human and *Drosophila* populations have found evidence of a percentage of synonymous mutations being under strong purifying selection (Keightley and Halligan, 2011; Lawrie et al. 2013; Ragsdale et al. 2018; Machado et al. 2020). Finally, some recent results report widespread selection acting on synonymous mutations (Shen et al. 2022a), however, the validity of these results is under debate (Shen et al. 2022b; Kruglyak et al. 2023). Nevertheless, the effect of selection on synonymous mutations on DFE inference of non-synonymous mutations remains unclear.

In this work, we address this knowledge gap by testing the robustness of one DFE inference method, Fit∂a∂i (Kim et al. 2017), to the presence of selection on synonymous mutations. First, we show that selection on synonymous mutations results in false evidence of population expansion events during the demographic inference step. Furthermore, we show that the selection acting on synonymous mutations generates clear differences in the DFE parameter estimates. However, these differences only result in significant mispredictions of the proportion of moderate and strongly deleterious mutations when selection on synonymous sites is widespread and very deleterious. Finally, we show that the misestimation of the demographic parameters is primarily responsible for the incorrect DFE inferences. This points toward a possible strategy to resolve the bias as well as a potential approach for detecting selection acting on synonymous mutations.

## Methods

### Simulations

To test the performance of Fit∂a∂i in the presence of selection on synonymous mutations, we simulated datasets with known selection parameters. Simulations were performed with the forward-in-time evolutionary simulation framework, SLiM 3 (Haller and Messer, 2019). A population with a constant size of 10000 individuals was simulated for 100000 burn-in generations, followed by an additional 1000 generations before sampling 100 haploid genomes. For each simulation, random exon and intron regions were generated across the total length of the simulated sequence with an average 1.4Mb in exonic sequence and 19.6Mb in intronic sequence. The mutation rate (*μ*), and recombination rate (*r)* were constant across the entire length of the sequence with *μ*=1.5e-8 per base pair per generation, and *r*=1e-8 per base position per generation, unless otherwise specified. Further simulation parameters are listed in Supplementary Table 1.

Two categories of mutations were simulated in the exonic regions: non-synonymous (NS) mutations and synonymous (S) mutations. The ratio of NS to S mutations in the simulation was 2.31:1 (Huber et al. 2017). We simulated a human-like nonsynonymous DFE (Kim et al. 2017) where the deleterious selection coefficients (*s*) for NS mutations were drawn from a gamma distribution with a scale parameter (in terms of 2*Ns*) of 706.899 and a shape parameter of 0.186, yielding a mean *s* of 0.013. The selection coefficients for the synonymous mutations depended on the model of selection on synonymous sites being tested. Under the simplest model, the constant model, all synonymous mutations were assigned the same deleterious selection coefficient. Three values of *s* were tested: a nearly neutral, *s*=1e-5 (*Ns* <1), a moderately deleterious, *s*=1e-4 (*Ns* ∼ 1), and a strongly deleterious, *s*=1e-3 (*Ns* > 10). Under the partial model, 22% of synonymous mutations were assigned the same selection coefficient. The three selection coefficients tested under the partial model: *s*=1e-5, *s*=1e-4 and *s*=1e-3. The remaining 78% of mutations were neutrally evolving and had *s*=0. The partial model reflects an attempt at a more biologically informed model for selection on synonymous sites based on estimates from *Drosophila melanogaster* (Lawrie et al. 2013). Accounting for the different synonymous selection coefficients, we conducted simulations for a total of six conditions with selection on synonymous mutations. We also generated a control condition where all synonymous mutations were neutral (*s*=0).

Each individual simulation run contained approximately 1.5Mb of coding sequence. To generate the data for a single replicate, we aggregated the results of 22 individual simulations, run in parallel to speed up computation. The results of each individual run were aggregated into a synonymous and non-synonymous site frequency spectrum (SFS). The respective SFS of 22 simulations were summed to obtain a single replicate synonymous and non-synonymous SFS. For each condition, we generated 20 replicates, where each replicate represents approximately 30Mb of coding sequence (1.5Mb per simulation ✕ 22 parallel simulations ∼ 30Mb).

### Recombination Simulations

To test the effects of linkage on our results, we varied the strength of recombination. We tested two possible elevated recombination rates, *r*=1e-7 and *r*=1e-6. We ran the simulations as described in the previous section, varying only the recombination parameter. For each of the seven possible conditions previously described (six conditions with selection on synonymous mutations plus a control condition), we generated 20 replicates with each one of the two elevated recombination rates.

### Demographic Inference

We used the software package ∂a∂i (Gutenkunst et al. 2009) to infer the demographic parameters that provided the best fit to the synonymous SFS obtained from each simulation replicate. We tested two possible demographic models: a one-epoch model representing a population with no size changes and is the true demography that was simulated, and a two-epoch model, which represents a single population size change at some point in the population’s history and fits two parameters.

We used a likelihood ratio test (LRT) to determine whether the two-epoch model fits the data significantly better than the one-epoch mode. The log-likelihood ratio test statistic between the null (one-epoch) and alternative (two-epoch) models can be asymptotically approximated under the null hypothesis by a *χ*^2^ distribution with 2 degrees of freedom (Wilks, 1938). A log-likelihood difference >3 represents a significance level of 0.05 and allows us to reject the one-epoch model in favor of the two epoch model for providing a statistically significant better fit to the data.

For each simulation replicate, we first tested 25 possible initial parameters in each round of model fitting. Initial parameters were tweaked until a good fit between the simulation SFS and the predicted SFS under the demographic model was achieved. Satisfactory fit was assessed through visual inspection of the true and predicted SFS for all demographic inferences for all replicates. We imposed an upper bound to the two epoch demographic parameters as follows: Upper bound for *v* (fold difference between current and ancestral population size) of 3000, upper bound for *T* (time of fold change in units of *2N_a_* generations) 500. A *v* value of 3000 led to small values of the population-scaled synonymous mutation rate, θ_s_∼1.5, which in turn corresponded to a highly biologically improvable ancestral population size of ∼3 individuals.

### DFE Inference

We used the software package Fit∂a∂i (Kim et al. 2017) to infer the parameters of the distribution of fitness effects (DFE) of the non-synonymous mutations. We conditioned the inference on the best fit demographic parameters obtained for each simulation replicate after following the inference procedure outlined above. We fit a gamma distributed DFE to the non-synonymous SFS, which was the functional form of the true DFE used to generate the data. For each replicate, we conducted 25 independent iterations of model fitting, varying the initial gamma distribution parameters. The parameters values having the maximum likelihood values of all the iterations were considered the best fit.

Fit∂a∂i reports the DFE in terms of the population-scaled selection coefficient, γ=2N_a_s, where N_a_ is the ancestral population size and *s* is the selection coefficient of the heterozygote. In order to obtain the DFE in terms of *s*, we first obtained the ancestral population size for a particular replicate from the synonymous population mutation rate estimated during the demographic inference step. Then, we used the fact that θ_s_ = 4N_a_μL_s_, where μ is the per-base-pair mutation rate and L_s_ is the length of possible synonymous sites, to obtain an estimate of the ancestral population size for a given replicate. The estimation of the population size is used to obtain the results in terms of *s* for that replicate using the formula for γ above. As well, we used it to compute the scale parameter in terms of the selection coefficient of the heterozygote, *s_dhet_* = *b / 2N_a_*. *b* represents the scale parameter in units of population-scaled selection strength directly obtained from Fit∂a∂i. Other parameters used in our study are outlined in Supplementary Table 1. For more details, see Kim et al. 2017.

The estimates of discretized DFE densities in each selective coefficient bin were obtained by drawing 10000 values from a gamma distribution for each set of best-fitting parameters. Then, we took the average across replicates for each model of selection on synonymous sites.

## Results

### Selection on synonymous mutations results in false evidence of population expansion

We first inferred the demographic history of each simulation replicate using the site frequency spectrum (SFS) of synonymous variants. In some parameter combinations, the synonymous variants were neutrally evolving, and in other conditions, they experienced negative selection as described in Methods. We fit two possible demographic models to the data: a one-epoch model, which corresponds to a constant population size throughout the entirety of the population’s history; and a two-epoch model, where a single population size change occurred at some point in the population’s history. Although we know that the two-epoch model is incorrect, as the data were simulated under a constant population size model, we selected these two models to represent the standard practice procedure expected when researchers perform DFE inference in a population of unknown demographic history (Boyko et al. 2008).

First, we tested a model where all synonymous mutations are deleterious with a single constant selection coefficient, which we will refer to as the “constant model”. As a control, we also performed a set of simulations with no selection on synonymous mutations. We observed a skew towards rare variants in the synonymous SFS when synonymous mutations experienced selection (Supplementary Figure 1 and 3). The skew increases with increasing strength of selection, as is to be expected due to the effect of purifying selection on the SFS. The shape of the non-synonymous SFS does not show significant changes when synonymous mutations experience selection (Supplementary Figure 3).

After demographic inference, the one epoch model is preferred in 14/20 replicates in our control simulations (*s*=0 on all synonymous sites). For the control simulations where the two-epoch model is preferred (6/20), only small expansions are predicted, with the fold difference between the current and ancestral population sizes is at most 1.1002 (Figure 1A, Supplementary Table 3). Inference of small expansions from simulated data without a true expansion has been previously reported (Kim et al. 2017; Messer and Petrov, 2013; Schrider et al. 2016) and is due to linked selection.

**Figure 1:**
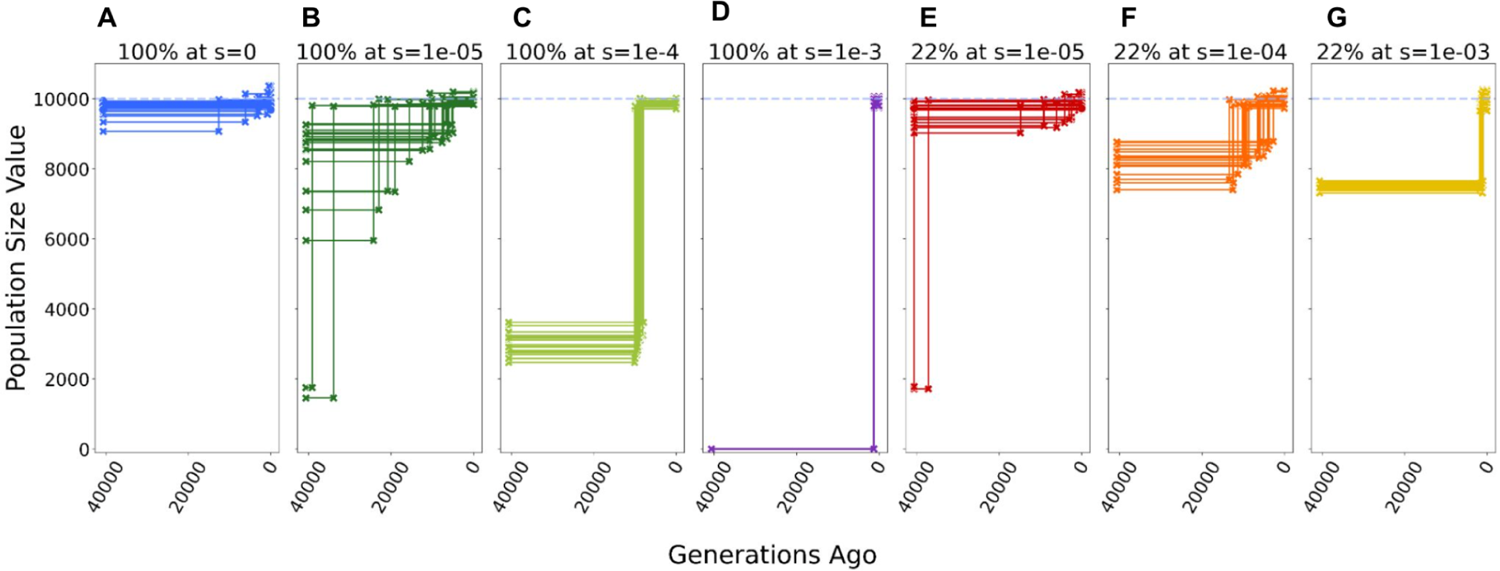
Inference of demography with varying degrees of selection on synonyms mutations. Inferred population size for each replicate under each model of selection on synonymous mutations. Each scenario includes 20 simulation replicates. When a Two Epoch model (one population size change at a specific time in the past) provides the best fit, the inferred time of the demographic event is indicated by a step in the plot between the ancestral population size and current population size. A horizontal line indicates data best described by a One Epoch (constant population size) model. Dashed blue line corresponds to the true population size in all simulations (*N*=10000). Inference in each replicate performed on a sample of 100 chromosomes. **A** Neutral model, with no selection on synonymous mutations (*s*=0). **B-D** Model with all synonymous mutations experience selection, with deleterious selection coefficients (s) ranging from 1e-5 to 1e-3. **E-G** Model where 22% of synonymous mutations experienced selection, with deleterious selection coefficients (*s*) ranging from 1e-3 to 1e-5. The remaining 78% of synonymous mutations are neutral (*s*=0).

In all replicates with constant selection on synonymous mutations, the two epoch model provides the best fit in all replicates, regardless of the strength of the synonymous selection coefficient. This prediction is consistent with the synonymous SFS being skewed towards rarer variants due to negative selection. With increasing strength of negative selection on synonymous mutations, the predicted difference between the ancestral and current population sizes becomes larger (Figure 1, B to D and E to G). Further, increasing selection led to more recent estimates of the expansion (Figure 1, B to D and E to G). In some replicates of the most extreme constant condition (*s*=1e-3, Figure 1D), we were unable to find a demographic model that would properly match the observed synonymous SFS. However, the best-fitting parameters represent the most extreme parameters allowed in our inference (See Methods). For example, the difference between the ancestral and current population size is estimated to be in the order of 3000-fold, which leads to ancestral population sizes in the order of ∼3 individuals (Figure 1D). Interestingly, when all synonymous mutations experience weak selection (*s*=1e-5, Figure 1B and E), the variance in the predicted timing and degree of the mispredicted population expansion increases. In some replicates, large expansions are predicted, larger even than those consistently predicted in the moderately deleterious constant condition (Compare Figure 1B to 1C).

Next, we examined the “partial model” where only 22% of synonymous mutations were under selection (See Methods). We again observe a skew towards rarer variants only in the synonymous SFS (Supplementary Figure 1). The skew becomes greater with increasing negative selection on the synonymous mutations, but overall the effect is less pronounced in the partial model than in the constant model. For the replicates with moderate and strong selection on synonymous mutations (*s*>1e-4), all show a significantly better fit for the population expansion model compared to the one-epoch model. For the weak selection condition (*s*=1e-5), 9/20 replicates report a one-epoch model as the best fit to the data. However, when the two epoch model provides the best fit, there is a great variability in the predicted size and timing of the expansion. These results are qualitatively similar to what we observed in the constant model, where weaker selection on the synonymous mutations increases the variance in the MLEs of the two-epoch model parameters.

In summary, in nearly all of the scenarios that include selection on synonymous mutations, demographic inference shows evidence of a population expansion, even though no population size change took place.

### Selection on synonymous mutations influences the inferred DFE for nonsynonymous mutations

Next, we interrogated how selection on synonymous mutations affects inference of the DFE of non-synonymous mutations. We performed DFE inference using the demographic parameters obtained from ∂a∂i for each simulation replicate paired with that simulation’s corresponding non-synonymous SFS. A gamma distribution, described by scale and shape parameters, was fit to the data using the software package Fit∂a∂i (Kim et al. 2017).

We observed systematic deviations from the true DFE parameters for non-synonymous mutations when selection was acting on synonymous mutations (Figure 2A, Supplementary Figure 6A) in most conditions. In general, the observed deviations between the true and inferred parameters can not be explained by the variance during the inference process: the difference between the true and inferred parameters in the control replicates where synonymous mutations do not experience selection (blue dots on Figure 1A) are systematically smaller than the difference between the true parameters and any inferred parameter of the replicates where synonymous mutations experience selection (compare distances within blue dots to distance between blue dots to any other colored dot). In the constant model, replicates show significant parameter estimate differences with respect to the true parameter values (dark green, bright green and purple dots in Figure 2A), with the deviations increasing in distance from the true values with increasing selective coefficient on synonymous mutations. This trend is extreme in the constant condition with the strongest selection on synonymous mutations (*s*=1e-03, purple dots in Figure 2A). When all synonymous mutations are strongly selected against, we were unable to fit a set of DFE parameters that predicted the observed non-synonymous SFS (See Supplementary Figure 5). In the partial model, replicates showed significant parameter estimation differences with respect to the true parameters in the conditions where selection on synonymous mutations was moderate or strong (orange and yellow dots in Figure 1A). The deviations increased, as in the constant model, with increasing selective coefficient on synonymous mutations. However, the condition where only 22% of synonymous mutations were nearly neutral was an exception (red dots in Figure 1A, Supplementary Figure 4A). In this particular condition of the partial model, the DFE parameter estimates are similar to the parameter estimates obtained in the control replicates (blue dots in Figure 1A, Supplementary Figure 4A), indicating that the low level of selective pressure on a fraction of the synonymous mutations did not severely impact our ability to estimate the true DFE parameters.

**Figure 2:**
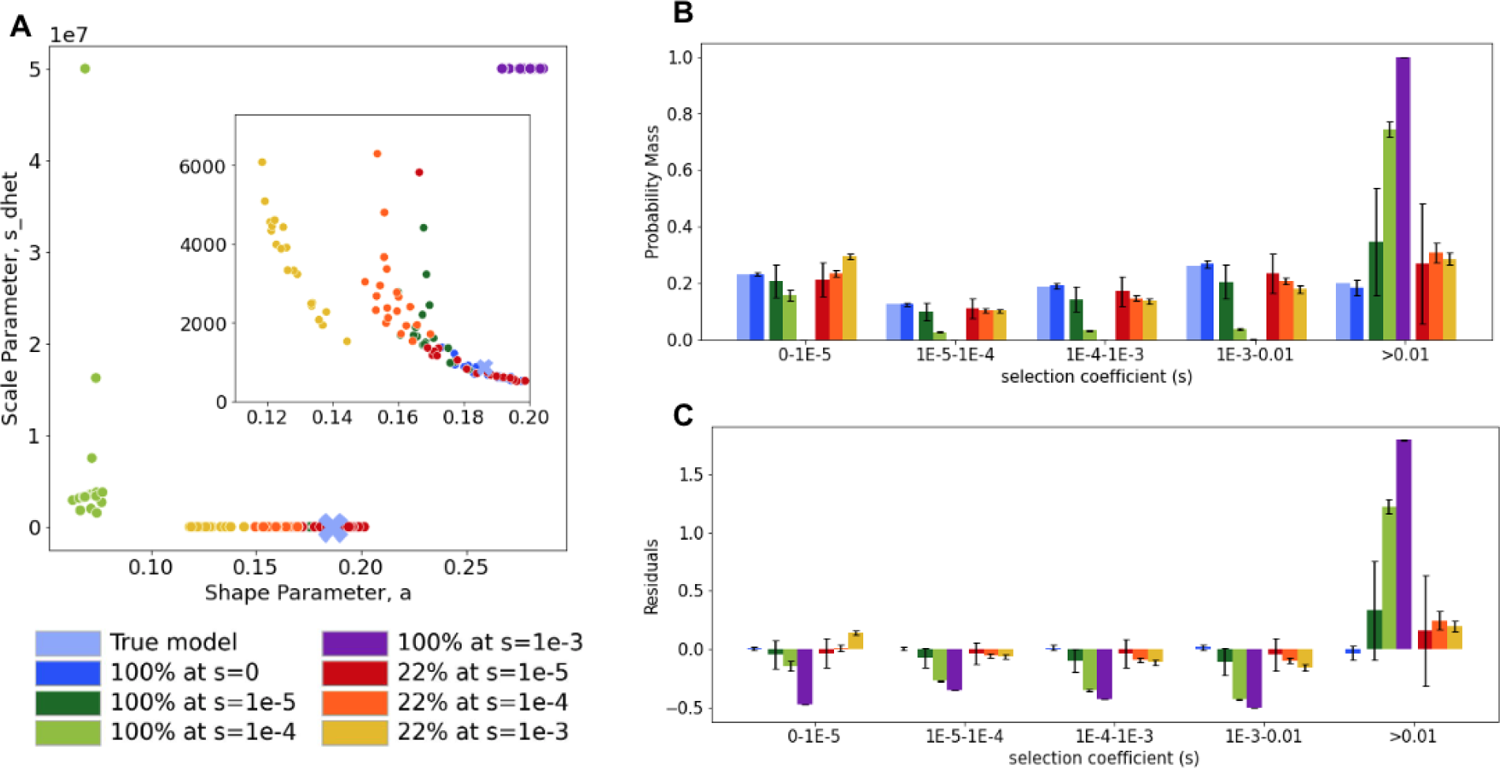
Inference of the distribution of fitness effects (DFE) for nonsynonymous mutations under different models of selection on synonymous mutations. **A** Inferred shape and scale parameters in a gamma DFE model for non-synonymous mutations from simulated data with distinct levels of selection on synonymous mutations. Each point represents an individual simulation replicate. Scale parameter, *s_dhet_*, represents the scale parameter in units of heterozygous selection strength. Insert zooms in on the lower-middle section of the plot. **B** Comparison of the discretized DFE for non-synonymous mutations between the true DFE (light blue) and the average inferred DFE for each model of selection on synonymous mutations. Bars represent an average of 20 replicates, error bars show standard deviations. DFE bins range from neutral (0-1E-5) and nearly neutral (1E-5-1E-4) to strongly deleterious (>0.01). **C** Standardized residuals of the probability mass in each DFE bin, obtained by subtracting the true probability mass (light blue in B) from the average for each condition, for each bin, divided by the square root of the true probability mass.

In the conditions with weak selection on synonymous mutations (both the partial and constant models with *s*=1e-05), we observed elevated variance in the inferred DFE parameters (dark green and red dots in Figure 2A and Supplementary Figure 4B). While most parameter estimates in both conditions cluster together, a few inferences report significantly elevated values of shape and scale (Supplementary Table 2, replicate 5 or 20 in the partial model, or replicate 12 and 15 in the constant model, Supplementary Figure 4B). These replicates correspond to cases where the inferred demography showed extreme population expansions. Since our ability to correctly infer the DFE parameters is conditional on controlling for demographic events, it follows that incorrect demographic parameters would impact our ability to estimate the DFE parameters.

Next, we checked whether inferences of the proportions of mutations with different selection coefficients were also affected given the deviations we encountered in the DFE parameters from conditions experiencing selection on synonymous mutations. To do this, we divided the DFE into 5 bins and found the probability mass in each bin from the MLE parameters of the gamma distribution (Figure 2B-C and Supplementary Figure 6B-C; see Methods). All models with selection on synonymous mutations showed at least slight deviations from the true proportion of mutations in each bin of the DFE. The degree of deviation from the true proportion of mutations in each bin of the DFE varied across models. For some scenarios, selection on synonymous mutations had little to no effect on the inference of the proportions of mutations in each bin of the DFE. For example, the partial conditions (red, orange and yellow bars in Figure 2B-C) and the constant conditions with weak selection (dark green bars in Figure 2B-C) showed little bias from the true DFE proportions. However, as the standard deviations are narrow for the partial conditions with moderate or strong selection (orange and yellow bars in 2B-C), the small bias is likely not due to simulation variance or statistical uncertainty. For other scenarios involving stronger selection on all synonymous mutations, the bias was more pronounced (light green and purple bars in Figure 2B-C). In these conditions, there is a large excess of highly deleterious non-synonymous mutations predicted (*s*>0.01). In the most extreme condition, where all synonymous mutations experience a coefficient of *s*=1e-3 (purple bars in Figure 2B-C) nearly all of the probability mass (99.8%) is located in the most deleterious category of the non-synonymous DFE. The large standard deviations observed in the categories experiencing weak selective coefficients on synonymous mutations (dark green and red in Figure 2B-C) are caused by the previously described variability in the inferred DFE parameters. In summary, the selective pressures on synonymous mutations can affect inferences of the DFE for nonsynonymous mutations.

### Decreasing linked selection through increasing recombination does not improve accuracy of DFE parameter estimates

We next explored which factors could have biased the parameter estimates of the DFE for nonsynonymous mutations when selection acted on synonymous mutations. Linkage among sites experiencing selection impacts patterns of variation across the genome (Murphy et al. 2022; Elyashiv et al. 2016; Charlesworth and Jensen, 2021; Cutter and Payseur, 2013; Lohmueller et al. 2011; Hernandez et al. 2011; Sella et al. 2009; Cai et al. 2009; Charlesworth et al. 1993; Fay and Wu, 2000). In particular, background selection affects our ability to infer the demographic history of a population (Johri et al. 2021; Pouyet et al. 2018; Schrider et al. 2016; Ewing and Jensen, 2015; Messer and Petrov, 2013; Comeron, 2014). Given the large number of synonymous mutations potentially experiencing negative selection in our simulations, we tested how linkage affects the demographic and DFE inferences.

To do this, we performed additional simulations with higher recombination rates to reduce linkage. By increasing the recombination rate, we expect to break haplotype blocks, and therefore unlink mutations. We selected two elevated recombination frequencies, one which represents a ∼10x increase in recombination rate with respect to the mutation rate (*r*=1e-7) and one that represents a ∼100x increase (*r*=1e-6). As a control, we first performed simulations with elevated recombination rates on a model without selection acting on synonymous mutations (Supplementary Figure 7). We observed similar accuracy in the inference of the DFE of nonsynonymous mutations regardless of recombination rate for the control simulations where synonymous mutations were neutral, indicating that recombination rate by itself does not impact DFE parameter inference. We did, however, observe an elevated number of simulations reporting a one-epoch model as the best fitting model with an increase in recombination (Supplementary Table 3).

Turning to the simulations with selection on synonymous mutations, we observed that the demographic inferences were closer to the true model with increased recombination in the partial model conditions. There was a greater number of replicates where the one-epoch was selected as the best-fitting model in the condition with weak partial selection on synonymous mutations (Supplementary Table 3). However, in the partial models with weak and moderate selection on synonymous mutations (red and orange lines in Supplementary Figure 8B) as well as in the constant model with week selection on synonymous mutations (dark green in Supplementary Figure 8A), there is an increase in the variance of the reported ancestral population size with increase in recombination rate. In the strong partial selection condition (yellow lines in Supplementary Figure 8B), the two-epoch model still provides the best fit to all replicates, but there is a consistent increase in the reported ancestral population size (*N_a_*) of the population (Supplementary Figure 9).

Increasing the recombination rate does not bring the inferred DFEs for nonsynonymous mutations closer to the underlying true DFE used in the simulations (Figure 3). When increasing recombination rates, we observed an increase in the proportions of mutations predicted to have strong selective coefficients (*s*>0.01) in all conditions, except under the partial condition where synonymous mutations are experiencing a strong selective coefficient. Here, we actually observed a slight decrease in the proportion of mutations that fall into the strongly selected-against bin (yellow bars in Figure 3B). This effect could be caused by the small increases observed in the reported ancestral population size (*N_a_*) of the population for this particular model only (Supplementary Figure 9)

**Figure 3:**
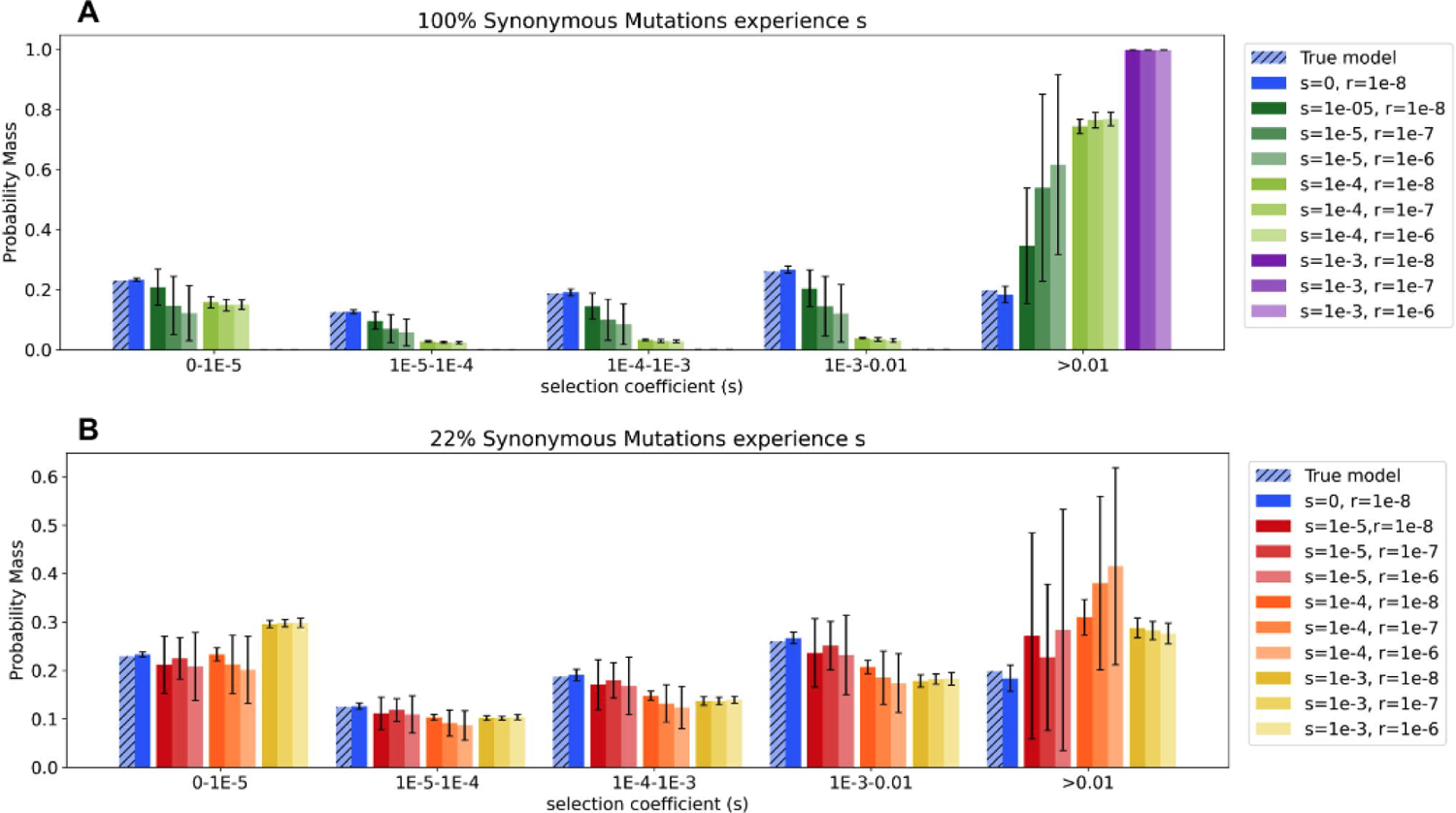
Inference of the Distribution of Fitness Effects (DFE) of non-synonymous mutations under varying strengths of recombination. Comparison of the discretized DFE for non-synonymous mutations between the true DFE and the average inferred DFE for each model of selection on synonymous mutations. Bars represent an average of 20 replicates, error bars show the standard deviations. DFE bins range from neutral (0-1E-5) and nearly neutral (1E-5-1E-4) to strongly deleterious (>0.01). Lighter colors indicate increasing recombination rate, *r*. **A** Discretized DFE for conditions where 100% of synonymous mutations experience selection. **B** Discretized DFE for conditions where 22% of synonymous mutations experience selection.

For the conditions with weak selection on synonymous mutations (dark green bars on Figure 3A and red bars on Figure 3B) as well as the partial model with moderate selection on synonymous mutations (orange bars in Figure 3B), there is a large variance in the proportion of mutations inferred in each bin of the DFE. This large variance corresponds to models where there is a large variance reported in the ancestral population size when a two-epoch model provides the best fit to the synonymous SFS (Supplementary Figure 8).

In summary, increasing the recombination rate does not improve our ability to infer the DFE when synonymous mutations are experiencing selection. We therefore conclude that linkage is not a major contributing factor to our inability to infer accurate DFE parameters under conditions experiencing selection on synonymous mutations.

### Using unlinked truly neutral variants improves DFE inference

Inserts zoom into points around true DFE parameters used to simulate the data, indicated by a blue cross. As we have shown, selection on synonymous mutations can confound inference of demographic history (Figure 1). Because the DFE inference conditions on the inferred demographic history of the population, we hypothesized that using the mis-inferred demographic parameters led to the biases in the inferences of the nonsynonymous DFE.

To test this idea, we assumed that there was a set of truly neutral variants that is known and can be used for demographic inference. These truly neutral variants could either be synonymous variants that have been validated to be truly neutrally evolving or some other set of noncoding polymorphisms in the genome. Importantly, we assumed that these neutral variants were unlinked from the nonsynonymous variants, so there is no need for them to be physically nearby the nonsynonymous variants. To simulate this scenario, we paired every non-synonymous SFS from each simulation replicate experiencing selection on synonymous sites with a set of demographic parameters inferred from our neutral simulations (Supplementary Figure 10). Since all our simulations share the same demographic history, the demographic parameters obtained from the neutral simulations represent an estimate of the demographic parameters we would expect to obtain if we had access to a set of known neutral sites.

When we conditioned our DFE inferences on demographic parameters obtained from known neutral variants, we were able to recover the true DFE parameters under all models of selection on synonymous mutations (Figure 4). This is reflected by the fact that the points all cluster more closely to the blue cross which denotes the true parameter values. Furthermore, the distribution of the shape and scale parameters from the inferences where demography was inferred from truly neutral sites are indistinguishable from the DFE parameter distributions obtained from our control simulations (Supplementary Figure 11).

**Figure 4:**
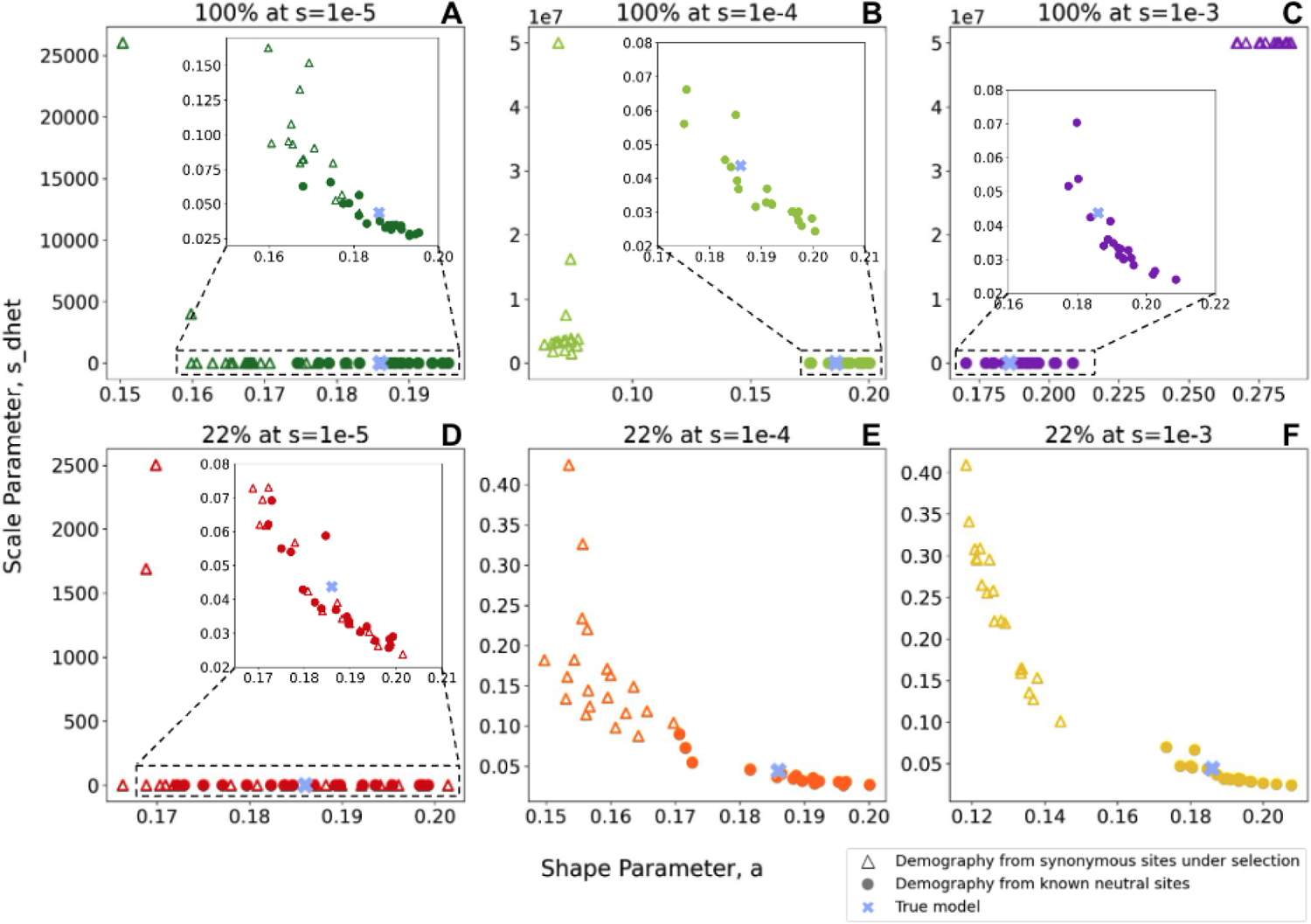
Comparison of inferences of the DFE of non-synonymous mutations with and without demography inferred from a known set of neutral variants. Inferred shape and scale parameters in a gamma DFE model for non-synonymous mutations from simulated data with distinct levels of selection on synonymous mutations. Each point represents an individual simulation replicate. Empty triangles represent parameter inferences generated using a demographic model inferred from a synonymous SFS experiencing selection, identical data as Figure 2A. Filled circles represent parameter inferences generated using a demographic model inferred from known neutral variants. Scale parameter, *s_dhet_*, represents the scale parameter in units of heterozygous selection strength. Each plot shows inferred parameters for a distinct model of synonymous selection: **(A-C)** constant model where all mutations are affected by selection; **(D-F)** partial model of selection, where 22% of the mutations are under selection.

The partial model with *s*=1e-5 is the only model where there is little difference between the parameters inferred using the standard pipeline versus the truly neutral demographic history (Figure 4D, insert). However, when using the standard pipeline in the partial model with *s*=1e-5, we encountered some replicates that reported significant differences with respect to the true DFE parameters (red triangles in the upper left corner of Figure 4D), which correspond to the cases with the best fitting demographic models corresponding to large expansions. When using demographic parameters obtained from known neutral sites on the partial model with *s*=1e-5, we do not observe any replicates with comparable differences with respect to the true DFE parameters (red circles in Figure 4D).

In summary, conditioning DFE inferences on demographic parameters inferred from a set of truly neutral variants allows us to recover the true DFE parameters for non-synonymous mutations, even when the truly neutral sites have independent genealogies (i.e. are unlinked) from the non-synonymous mutations. This result provides further understanding as to how selection on synonymous mutations confounds inference of the nonsynonymous DFE. Specifically, selection on synonymous mutations leads to misspecification of the demographic history when inferring the DFE for non-synonymous mutations.

## Discussion

In this work, we investigated the effects of selection on synonymous mutations on site-frequency spectrum-based methods for estimating demographic history and distribution of fitness effects (DFE) of non-synonymous mutations. We made use of forward-in-time evolutionary simulations to show the impacts of varying levels of selective pressure on synonymous mutations when conducting evolutionary inferences. As previously suggested (Ragsdale et al. 2018), we found that inferring demographic parameters from synonymous variants experiencing selection leads to false estimates of population expansion. The inferred fold expansion (ratio between the current and ancestral population size) increased with increasing negative selection on synonymous mutations. We then proceeded to infer the DFE of non-synonymous mutations from data simulated with synonymous mutations experiencing selection. We found systematic deviations in the inferred DFE parameters under many selection scenarios, except when selection is weak (*s*=1e-5) and acts only on a proportion of synonymous mutations (22%). These deviations sometimes resulted in overestimating the proportion of highly deleterious nonsynonymous mutations, particularly when selection is strong and is acting on all synonymous mutations. We showed that this effect is primarily mediated by conditioning the DFE inferences on incorrect demographic parameter estimates obtained from synonymous variation.

The excess of inferred strongly deleterious mutations can be understood by realizing the interplay between the demographic correction and the shape of the non-synonymous SFS. Due to the purging effect of purifying selection acting on non-synonymous mutations, the synonymous SFS is skewed towards rare variants. However, an excess of rare variants in the SFS can also be caused by a recent population expansion. Since the purifying selection on synonymous variants is not accounted for, ∂a∂i interprets the excess of rare variants as a demographic effect, in particular, an expansion having impacted the population. Demographic events impact synonymous and non-synonymous variation equally, therefore, we expect an excess of rare variants in the non-synonymous SFS as well. However, there isn’t an excess of rare variation in the non-synonymous SFS since no expansion event actually occurred. To compensate for the expectation of this ghost expansion, Fit∂a∂i assumes that the missing rare variants in the non-synonymous SFS are due to strong negative selection, and therefore reports DFE parameters that include stronger selection acting on the non-synonymous variation than was actually the case.

The main implication of our results is that studies reporting demographic and DFE inferences based on ∂a∂i and Fit∂a∂i may report inaccurate results if the selective pressures on synonymous sites are not accounted for. The predominant effect of selection on synonymous sites results in a consistent overestimation of a population expansion even when the selective coefficient on the synonymous mutations is weak, with *Ns* < 1(Figure 1B and E). The extent of the overestimation will be dependent on the strength of selection and the amount of synonymous variation experiencing selective pressures. Under the current evidence for selective pressures on synonymous mutations (Lawrie et al. 2013, Keightley and Halligan, 2011), which most closely resembles our data simulated under a partial synonymous DFE with *s*=1e-4, estimates of the fold-expansion of a population could be overestimated by 20% on average (Figure 1F). The impact on the estimated DFE parameters and the proportion of mutations predicted to be in each bin of the DFE is less severe when only a fraction of synonymous mutations experience selection (Figure 2), suggesting some robustness to unaccounted for selection on synonymous mutations. Further, our work indicates that previous inferences of the DFE for nonsynonymous mutations (Kim et al. 2017; Keightley and

Eyre-Walker, 2007; Tataru and Bataillon, 2019; Li et al. 2010; Huang et al. 2021; Castellano et al. 2019; Galtier, 2016, Chen et al. 2017) are not grossly mis-estimated. However, even under the current best estimates of selection effects on synonymous mutations (Figure 2B, orange bars), the proportion of strongly deleterious mutations (*s*>0.01) is consistently over-estimated. For example, in work, ∼30% of mutations are inferred to be strongly deleterious, when in truth only ∼18% are strongly deleterious. This difference might have practical relevance when the DFE is used for decision making in conservation efforts of endangered populations, where the amount of strongly deleterious variation present in the population is one of the main determinants of extinction risk due to inbreeding depression (Kyriazis et al. 2021).

Our study points to a potential solution to inferring demography and the DFE in the presence of selection on synonymous mutations. Specifically, the main contributor to the inaccuracy in the inferred DFE parameters of non-synonymous mutations is the use of incorrect demographic parameters (Figure 4). We show that using a demographic model inferred from known neutral variants improved the fit and accuracy of the inferred DFE parameters. Hence, one solution would be to use truly neutrally evolving sites for demographic inference instead of synonymous mutations. For example, a potential approach consists of using short intronic variation to obtain a neutral SFS from which the demographic history of the population can be inferred. Previous studies in Drosophila have shown that variants in short introns, defined as introns less than 65bp, experience little selective pressures in *Drosophila* at the level of polymorphism variants, specially between the 8-30bp positions (Parsch et al. 2010, Clemente and Vogl, 2012). Several studies have made use of short introns as a neutral reference, but a systematic approach has not yet been developed (Racimo et al. 2014; Eyre-Walker and Keightley, 2009; Lawrie et al. 2013). Another possible source of known neutral variation consists of ancestral transposable elements, although their use has only been proposed in mammals (Lunter et al. 2006). Finally, some studies have also used intragenic variants from putatively neutral regions as a reference (Ragsdale et al. 2018; Gazave et al. 2014).

One main advantage to using synonymous variants for demographic inference is that they are interdigitated with the nonsynonymous mutations and thus provide an approximate control for the effects of linked selection, which affects both synonymous and nonsynonymous mutations (Kim et al. 2017). If the short introns used for demographic inference are chosen to be located nearby the exons, then they may still be sufficiently linked to the nonsynonymous mutations to account for the background selection effect, which would represent an advantage over using intragenic variants or transposable elements. However, at least under average human recombination rates (*r*=1e-8) in out simulated data, the degree of background selection was not sufficient to appreciably bias our inferences of the nonsynonymous DFE even when the neutral control variants to infer demography were unlinked to the nonsynonymous mutations (Figure 4).

If one has access to a known set of neutrally evolving sites, one potential test for selective pressure acting on synonymous mutations could be to compare the demographic parameters predicted from a set of neutrally evolving variants to the parameters estimated from synonymous variation, using ∂a∂i. If the parameters differ significantly, that would suggest that selection might be impacting the synonymous variation in the dataset.

Interestingly, we were unable to find a well-fitting demographic model for our data under conditions of pervasive and strong selection on all synonymous mutations. In particular, the inference reached the biologically informed parameter bounds, which were set to an expansion that was nearly impossible. This result indicates another possible check for selection on synonymous mutations: If the inference of demographic parameters from a natural population is proving difficult and leading to biologically nonsensical expansions, researchers should consider the possibility that there might be pervasive selection acting on synonymous mutations. In those scenarios, we would strongly recommend seeking an alternative source of neutrally evolving sites.

Our study has some limitations. We only modeled two very simple DFEs for synonymous mutations: a uniform DFE across all synonymous mutations and a partial DFE, where 22% of synonymous mutations have the same selection coefficient and the rest are neutral. While it has been established that the DFE of non-synonymous sites is well represented by a gamma distribution (Eyre-Walker et al. 2007; Kim et al. 2017; Wade et al. 2023), the shape of the DFE of synonymous sites has not been well characterized. However, some studies have shown evidence that at least a portion of synonymous sites are under moderate (*Ns*>1) to strong (*Ns*>10) selection in both *Drosophila* (Lawrie et al. 2013) and humans (Keightley and Halligan, 2011). Both studies report a similar proportion (∼20%) of synonymous sites under selection. Our simulations are meant to provide a benchmark for the effects of selection on synonymous sites in our current inference methods, and therefore we decided to include a partial DFE to reflect the most up-to-date knowledge on the possible distribution of selective forces in synonymous mutations. However, more research is needed to better understand the distribution of fitness effects of synonymous sites which will in turn allow us to better characterize the impacts of this selective pressure on our inference methods.

Our work tested how violating the long-held assumption that synonymous mutations are neutrally evolving impacts the ability to infer demographic histories and DFE parameters of non-synonymous mutations from genetic variation data. Reassuringly, our study suggests current DFE parameter estimates that make use of synonymous variants as part of their inference procedure are not grossly misestimated, but careful consideration is required when estimating demographic histories from synonymous mutations. However, more research is required to characterize the extent to which synonymous mutations experience selection.

## Acknowledgements

We thank members of the Lohmueller and Garud labs for their helpful input throughout the project. The authors would like particularly thank Jonathan C. Mah for his assistance with the DFE inference using Fit∂a∂i and suggestions on the manuscript and Chenlu Di for helpful comments on the manuscript. This work was supported by the National Institutes of Health grant R35GM119856 to K.E.L.

## Data Availability

Scripts for simulations and inferences using ∂a∂i and Fit∂a∂i located at https://github.com/amzurita/Synonymous_Selection_Project/tree/main

