## Supplementary Information for "The impact of non-neutral synonymous mutations when inferring selection on non-synonymous mutations"

### Supplementary Tables

***Supplementary Table 1: Simulation Parameters.***
Population genetic parameters used to run forward-in-time simulations. The parameters are as follows: $n_c$, number of haploid chromosomes sampled to obtain the SFS of a particular simulation run; $N_a$, ancestral population size in diploids; $\mu$, mutation rate per base pair per generation; $r$, recombination rate per base position per generation; $L_t$ total length simulated per simulation run; $L_e$, average exonic length simulated per simulation run, with standard error. The length is reported as an average due to the exon/intron pairs being randomly generated for each simulation run. $L_{ns}/L_s$ is the ratio of possible nonsynonymous to synonymous mutations, obtained from Huber et al. 2016. $L_{er}$ is the average exonic length per replicate, after aggregating 22 simulations.

| $n_c$ | $N_a$ | $\mu$ | $r$ | $L_t$ | $L_e$ | $L_{ns}/L_s$ | $L_{er}$ |
|---|---|---|---|---|---|---|---|
| 100 | 10000 | 1.5e-8 | 1e-8 | 1e8 | 1463488 +/- 1585 | 2.31 | 32206526 +/- 7102 |

***Supplementary Table 2: Demographic and DFE inference results.*** Results for each replicate under different models of selection on synonymous sites. Columns are as follows: Condition indicates the model of selection on synonymous sites in that particular simulation replicate. Replicate_ID indicates the replicate number. Syn_Theta indicates the population-scaled synonymous mutation rate, $\theta_s$. Na indicates the ancestral population size for that particular replicate, calculated from $\theta_s$. Best_Model indicates whether the one epoch or two epoch model provided a best fit to the data. Nu and T are the fold change with respect to the ancestral population size and the time of that change in units of $2N_a$ generations, respectively. If a One Epoch model provides the best fit to the data, Nu is 1 and T is 0. Shape_Param and Scale_Param are the shape (*a*) and scale (*b*) parameters inferred for the gamma DFE of non-synonymous mutations in that simulation replicate. S_dhet indicates the scale parameter in units of heterozygous selection strength, computed from the scale parameter (See Methods).

***Supplementary Table 3: Comparison of the one and two epoch model fits across simulation replicates.*** The table lists the number of replicates for which a one epoch or two epoch model provided the best fit to the data. First column indicates the model of selection on synonymous sites used to simulate the replicates. Second column indicates the recombination rate used in the simulation replicates. For the remainder of the replicates simulated with selection acting on synonymous sites, which are not listed here, a two-epoch model always provided the best fit to the data.

| Model of selection on synonymous sites | Recombination rate | One epoch demographic model is best | Two epoch demographic model is best |
|---|---|---|---|
| 100% with *s*=0 | 1e-8 | 14 | 6 |
| 100% with *s*=0 | 1e-7 | 16 | 4 |
| 100% with *s*=0 | 1e-6 | 19 | 1 |
| 22% with *s*=1e-5 | 1e-8 | 9 | 11 |
| 22% with *s*=1e-5 | 1e-7 | 10 | 10 |
| 22% with *s*=1e-5 | 1e-6 | 14 | 6 |

### Supplementary Figures

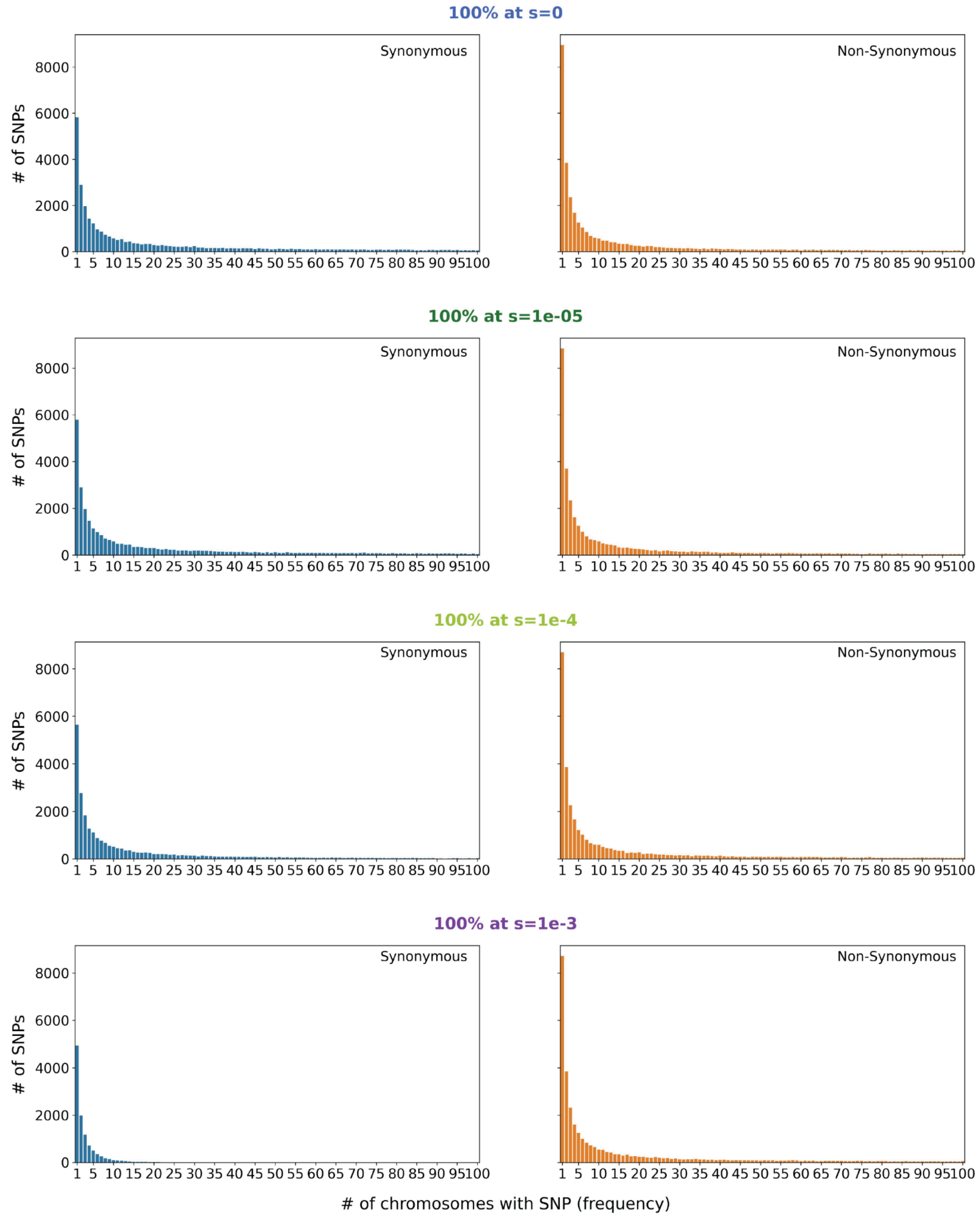

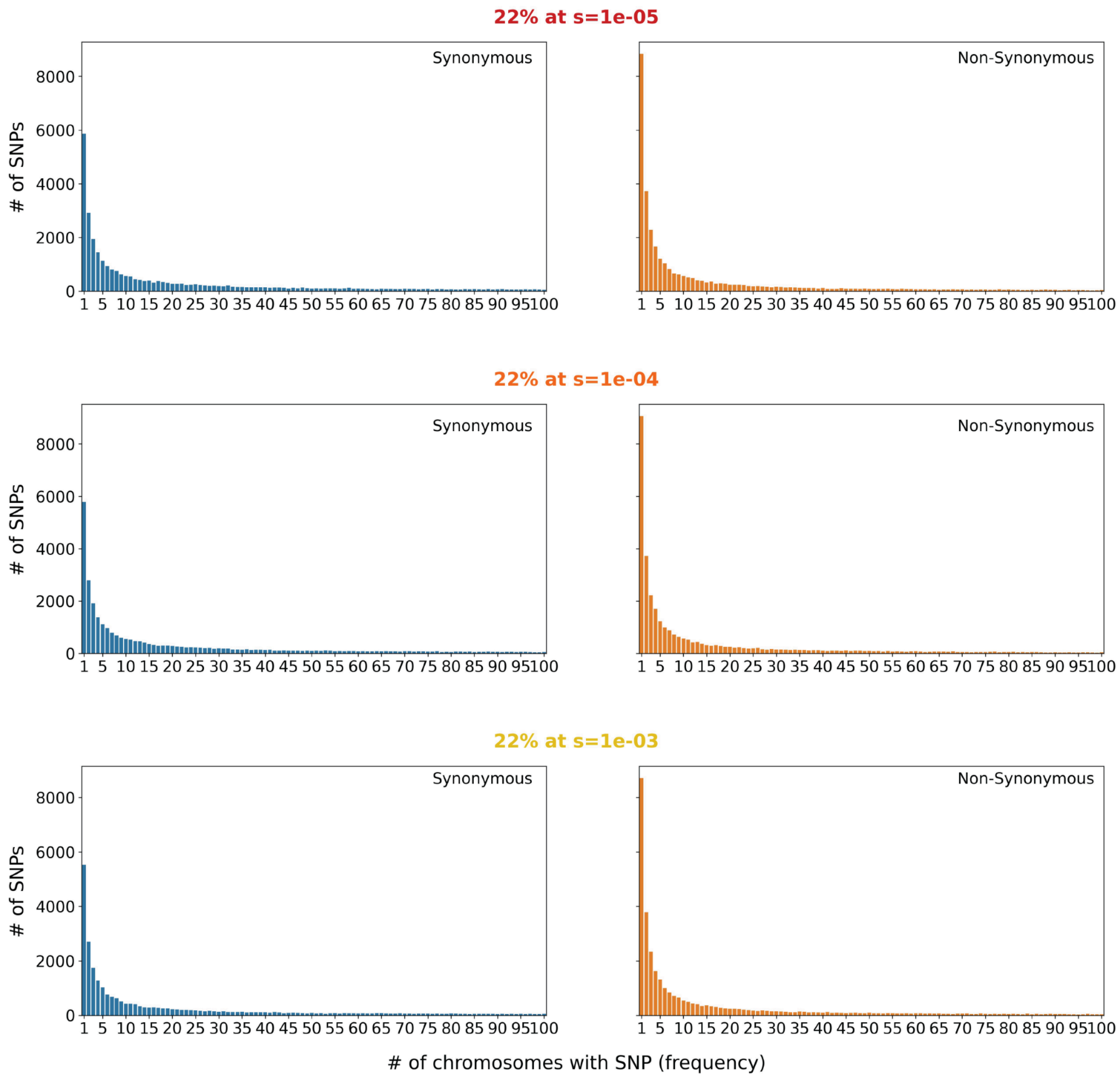


<u>**Supplementary Figure 1:**</u> **Site Frequency Spectra (SFS) for synonymous and non-synonymous variants obtained from a single simulation replicate**. Each row corresponds to a replicate from a specific model of selection on synonymous sites, as indicated by the title. The SFSs are obtained from a random sample of 100 haploid individuals collected at the end of the simulation. The x-axis represents the number of chromosomes in the sample with the particular variant. The y-axis shows the number of single nucleotide polymorphisms (SNPs) found at a particular count in the sample. For example, the 1 bin represents the number of variants that are only found in a single chromosome (singletons).

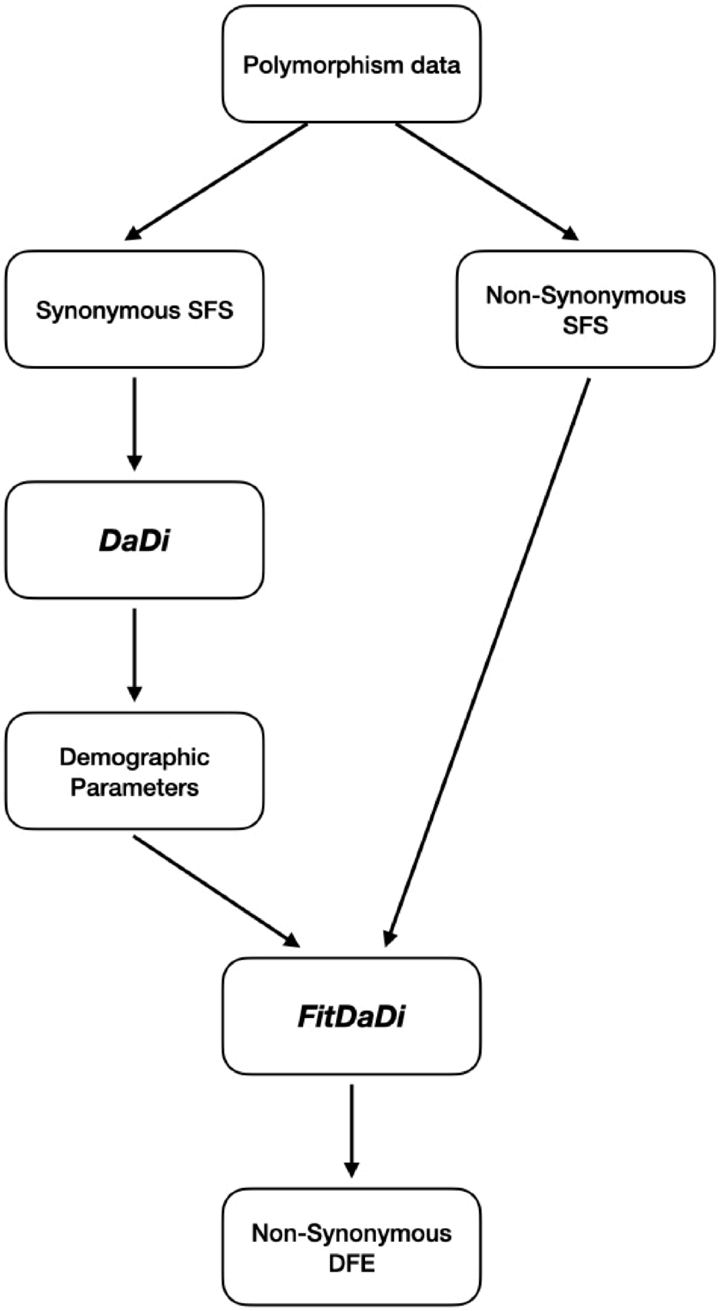


<u>Supplementary Figure 2:</u> **Schematic of the workflow used to obtain the parameters of the distribution of fitness effects of non-synonymous mutations from polymorphism data**. The bolded and italicized words represent specific software: ∂a∂i (Gutenkunst et al. 2009) is a demographic inference software that uses a diffusion approximation approach to compute the expected SFS under a particular demographic model. Fit∂a∂i (Kim et al. 2017) is a software package that does rapid inference of DFE parameters of new mutations.

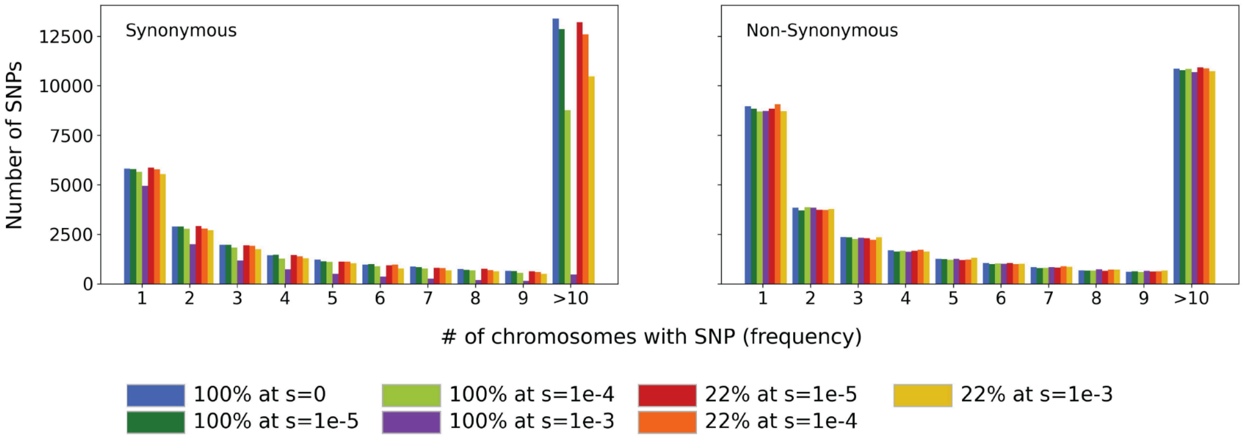


**<u>Supplementary Figure 3: </u>Comparison of SFS for synonymous and non-synonymous variants obtained from a single simulation replicate across models of selection on synonymous mutations**. The color of the bar indicates the specific model used in the simulation. The x-axis represents the number of chromosomes in the sample with the particular variant. Variants that are present in more than 10 individuals are summed and shown in the >10 bin, to make the visual comparison easier. The y-axis shows the number of single nucleotide polymorphisms( SNPs) found at a particular count in the sample

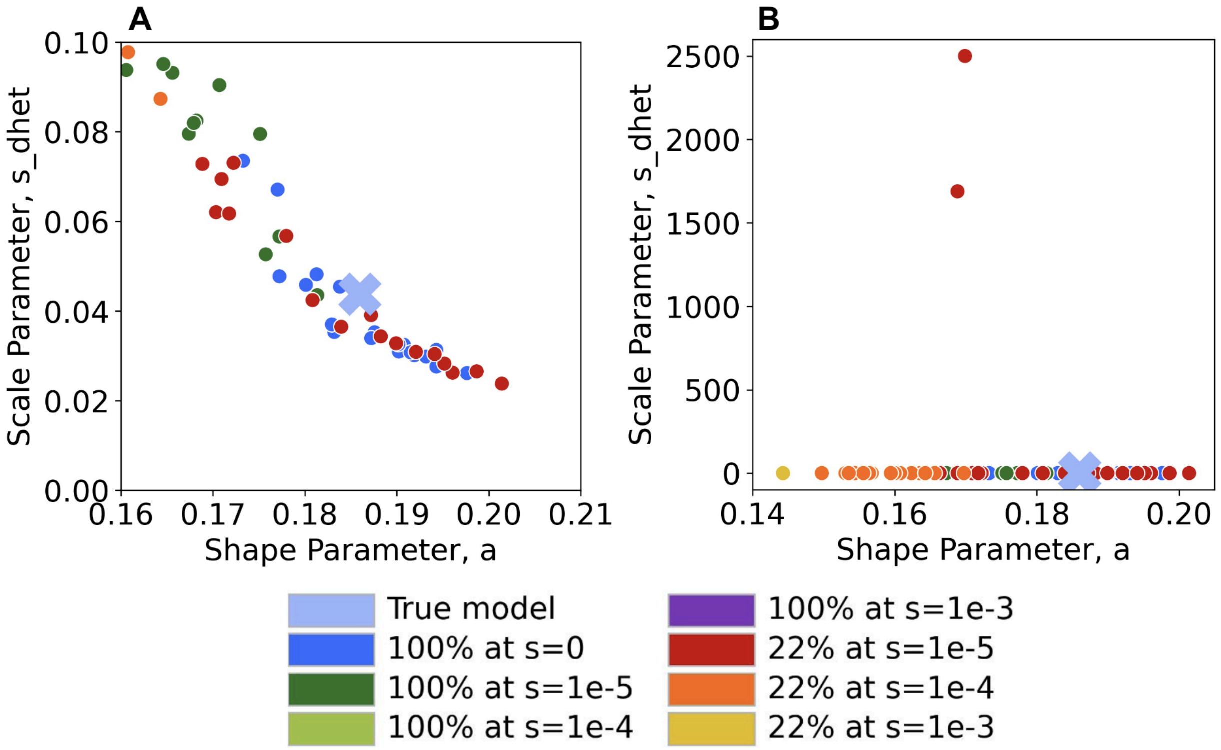


**Supplementary Figure 4: Inferred shape and scale parameters in a gamma DFE model for non-synonymous mutations from simulated data with distinct levels of selection on synonymous mutations**. Each point represents an individual simulation replicate. Scale parameter, $s_{dhet}$, represents the scale parameter in units of heterozygous selection strength. **A** Zoom into Figure 2 area around the true DFE parameters. $s_{dhet}$ limits range from 0 to 0.1. **B** Zoom into the lower area of Figure 2. $s_{dhet}$ ranges from 0 to 2600. Notice the y axis on both plots is only a small percentage of the y axis range in Figure 2.

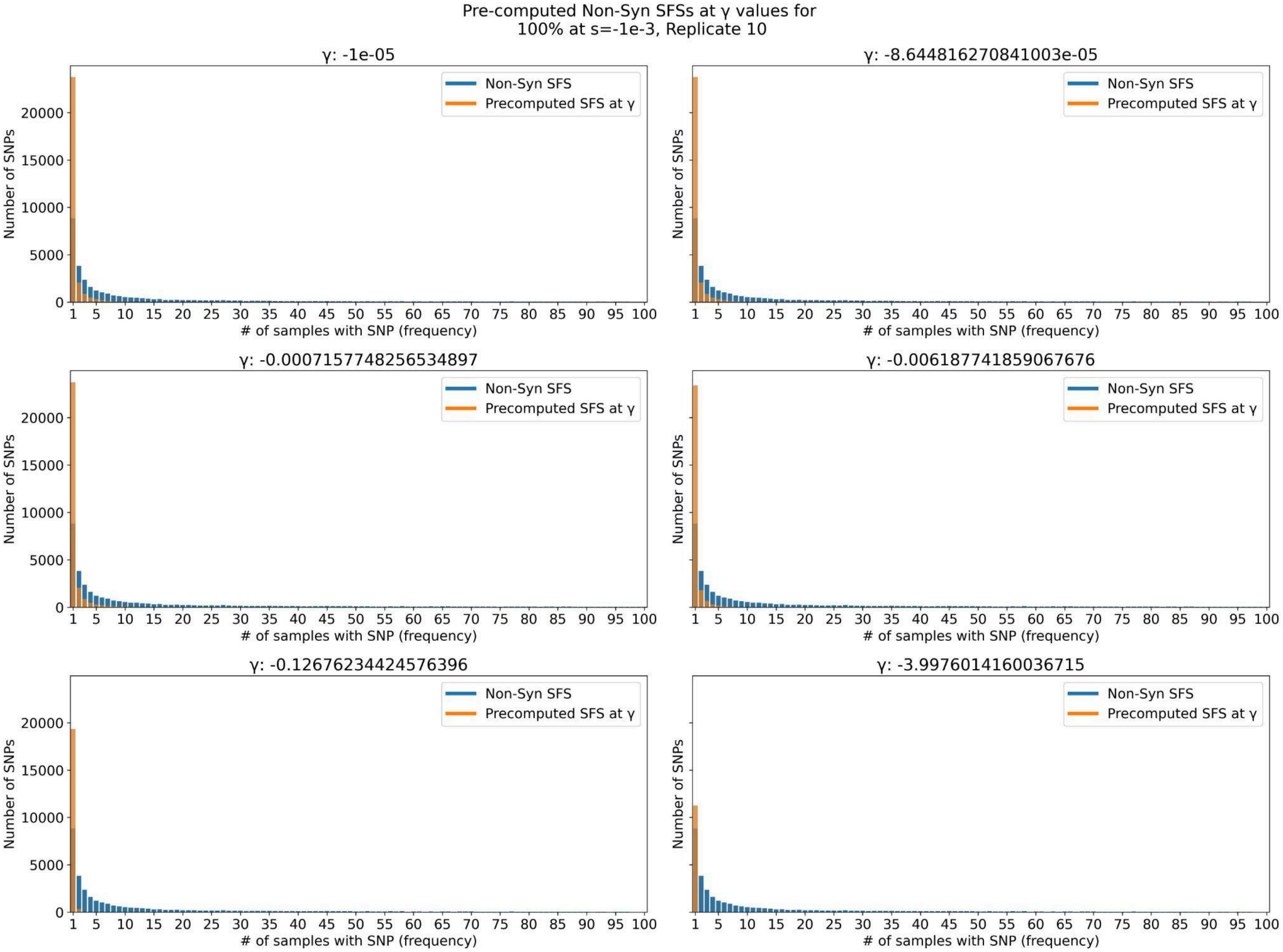


<u>**Supplementary Figure 5:**</u> **Pre-computed SFSs under γ values along the entire γ range for a replicate under the most extreme case of selection on synonymous sites, where 100% of mutations have *s*=1e-3**. Fit∂a∂i pre-computes the expected non-synonymous SFS under γ values ranging from 1e-05 to 2*$N_a$*0.5, where 0.5 is the most extreme selection coefficient possible since γ is the population-scaled selection coefficient of the heterozygote. The computation takes into account the demographic model. This is a representative replicate of the model where 100% of mutations have s=1e-3. None of the pre-computed SFSs can accurately represent the real non-syn SFS since there are no SNPs found in the common variants (frequency > 15) for the pre-computed SFSs (orange bars), but the real non-synonymous SFS has SNPs at common frequencies (blue bars).

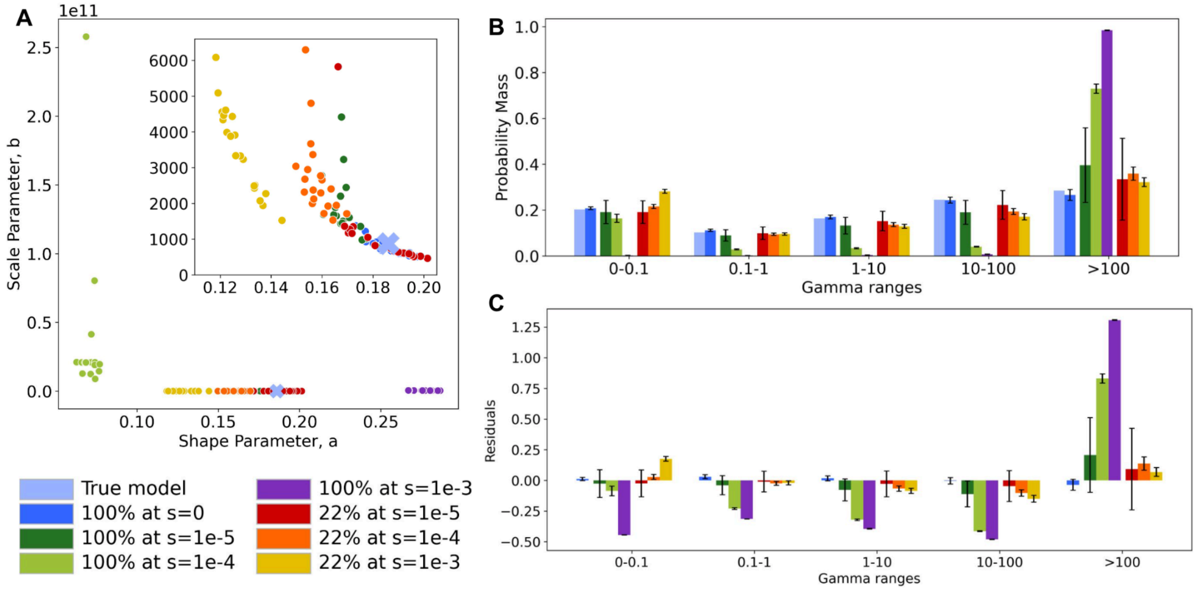


**<u>Supplementary Figure 6:</u> Inference of the distribution of fitness effects (DFE) for nonsynonymous mutations under different models of selection on synonymous mutations based on the population-scaled selection coefficient (γ).** Results from Figure 2 were obtained by scaling the results in terms of γ by the ancestral population size (see Methods) **A** Inferred shape and scale parameters in a gamma DFE model for non-synonymous mutations from simulated data with distinct levels of selection on synonymous mutations. Each point represents an individual simulation replicate. Scale parameter, *b*, represents the scale parameter in units of population-scaled selection strength. *b* relates to $s_{dhet}$ through scaling by $2N_a$, $s_{dhet} = b / 2N_a$. Insert zooms in on the lower-left section of the plot. **B** Comparison of the discretized DFE for non-synonymous mutations between the true DFE (light blue) and the average inferred DFE for each model of selection on synonymous mutations. Bars represent an average of 20 replicates, error bars show standard deviation. DFE bins range from neutral s (0-1) to strongly deleterious (>100). **C** Standardized residuals of the probability mass in each DFE bin, obtained by subtracting the true probability mass (light blue in B) from the average for each condition, for each bin, divided by the square root of the true probability mass.

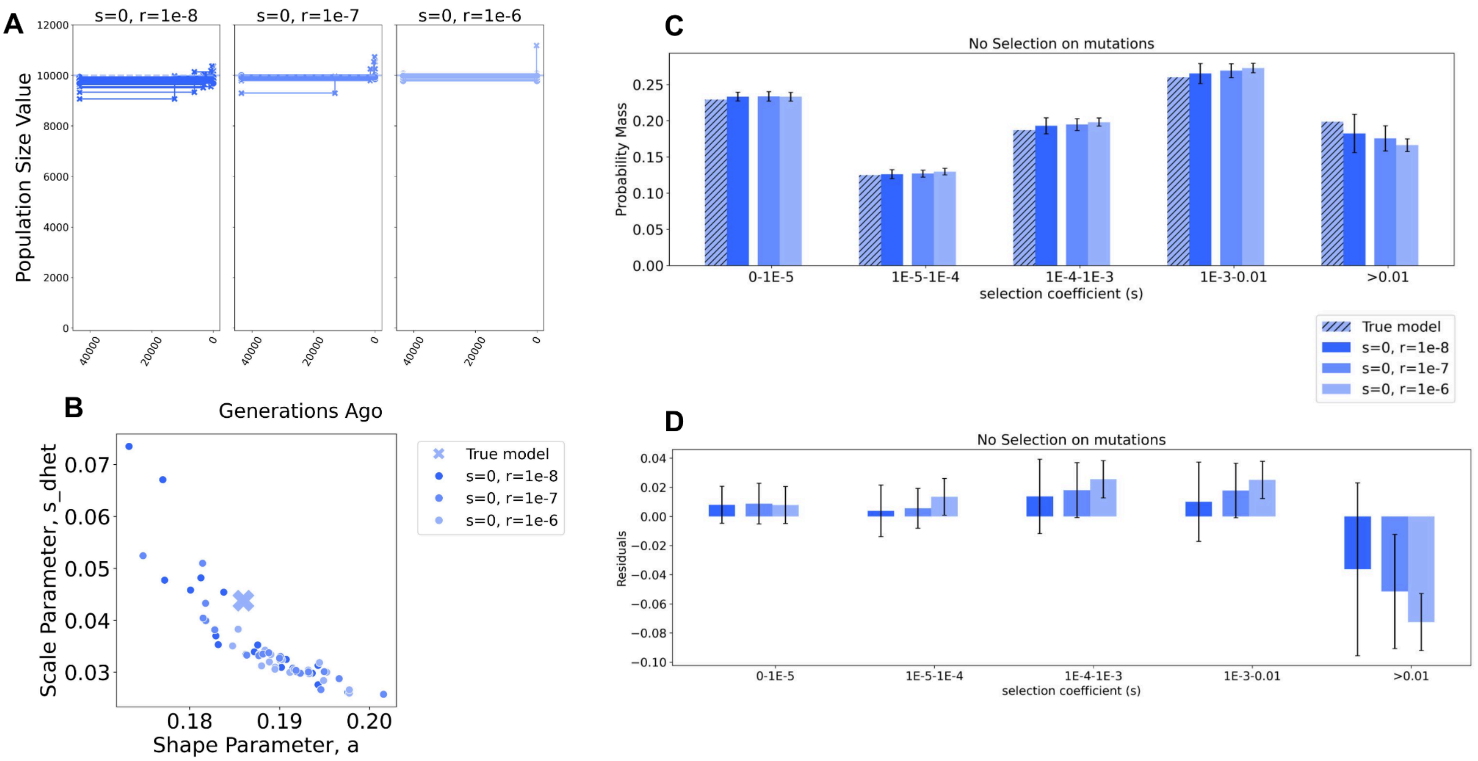


**<u>Supplementary Figure 7: </u>Inference of demographic and DFE parameters for the neutral model of selection on synonymous mutations when varying the rate of recombination (r)**. All replicates simulated with *s*=0 on synonymous mutations. Different shades of blue represent varying degrees of recombination. **A** Inferred population size for each replicate under each recombination rate tested. **B** Inferred shape and scale parameters in a gamma DFE model for non-synonymous mutations from simulated data with distinct levels of recombination. Each point represents an individual simulation replicate. Scale parameter, $s_{dhet}$, represents the scale parameter in units of heterozygous selection strength. **C** Comparison of the discretized DFE for non-synonymous mutations between the true DFE (striped blue) and the average inferred DFE. Bars represent an average of 20 replicates, error bars show standard deviations. DFE bins range from neutral selection values (0-1E-5) and nearly neutral (1E-5-1E-4) to strongly deleterious (>0.01). **D** Standardized residuals of the probability mass in each DFE bin, obtained by subtracting the true probability mass (striped blue in C) from the average for each condition, for each bin, divided by the square root of the true probability mass.

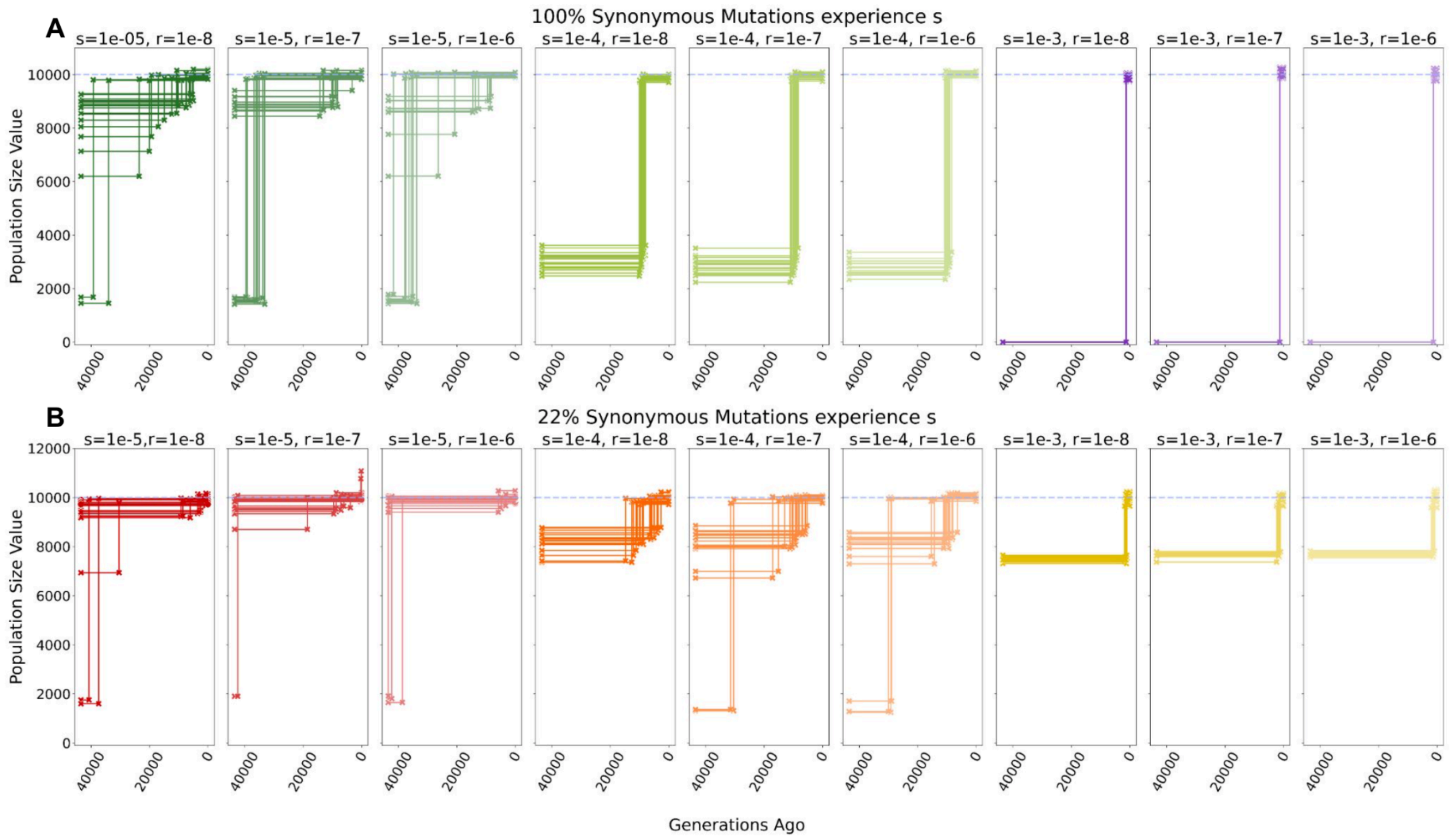


__Supplementary Figure 8:__ **Inference of demography with varying degrees of selection on synonyms mutations and recombination rates.** Inferred population size for each replicate under each model of selection on synonymous mutations and replication rate. Each scenario includes 20 simulation replicates. When a Two Epoch model (one population size change at a specific time in the past) provides the best fit, the inferred time of the demographic event is indicated by a step in the plot between the ancestral population size and current population size. A horizontal line indicates data best described by a One Epoch (constant population size) model. Dashed blue line corresponds to the true population size in all simulations (*N*=10000). Inference in each replicate performed on a sample of 100 chromosomes. **A** Demographic inferences for models of selection on synonymous sites where 100% of synonymous mutations experience selection. Specific selection coefficient and recombination rate of replicates indicated by *s* and *r*, respectively, at the top of each plot. **B** Demographic inferences for models of selection on synonymous sites where 22% of synonymous mutations experience the selection coefficient. Specific selection coefficients and recombination rates of replicates are indicated by *s* and *r*, respectively, at the top of each plot.

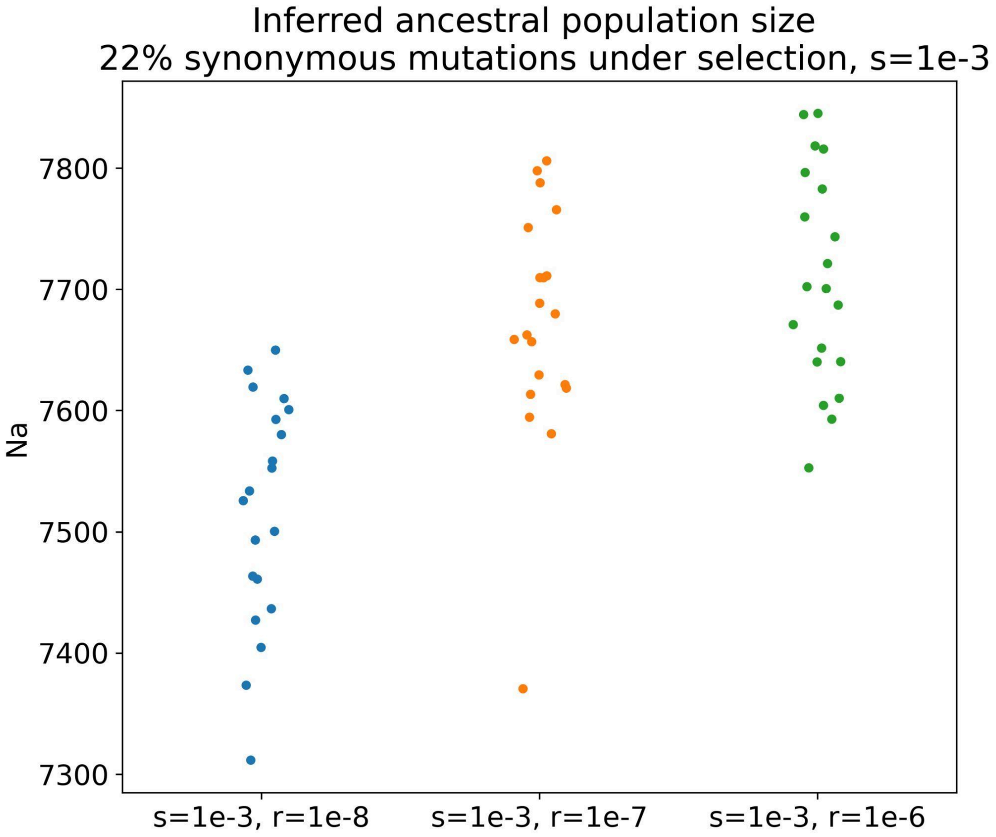


**Supplementary Figure 9: Inferred ancestral population size for replicates with 22% of synonymous mutations experiencing a selection coefficient of *s*=1e-3 with increasing recombination rate, *r*.** Each dot represents an individual replicate.

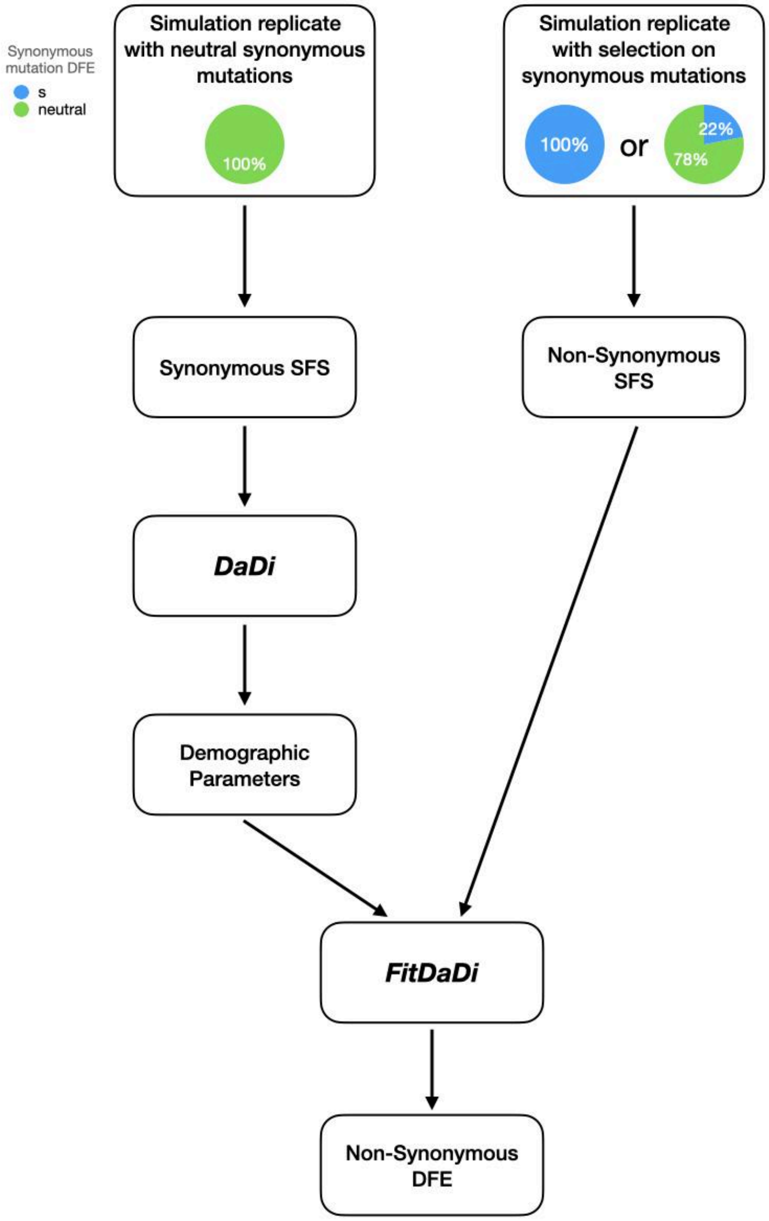


**<u>Supplementary Figure 10:</u> Schematic of the procedure followed to test the usefulness of a set of known unlinked neutral variants when inferring the DFE of non-synonymous mutations.** The non-synonymous SFS of a single replicate from a simulation where synonymous sites experience selection (right side) is paired with the demographic parameters inferred from a simulation replicate without selection acting on the synonymous mutations (left side). Fit∂a∂i performs the inference of DFE of non-synonymous mutations conditioned on the demographic parameters inferred from known neutral sites. For each of the 20 replicates simulated under each model of selection on synonymous sites, we paired the replicate with a set of demographic parameters inferred from a simulation where synonymous sites were entirely neutral. We conditioned the DFE inference on the demographic parameters. For example, in replicate 1 of the constant simulation with *s*=1e-5, we conditioned the DFE inference on the demographic parameters inferred from replicate 1 of the control simulations, where *s*=0 on all synonymous sites. Non-synonymous mutations in all simulations have the same DFE (See Methods).

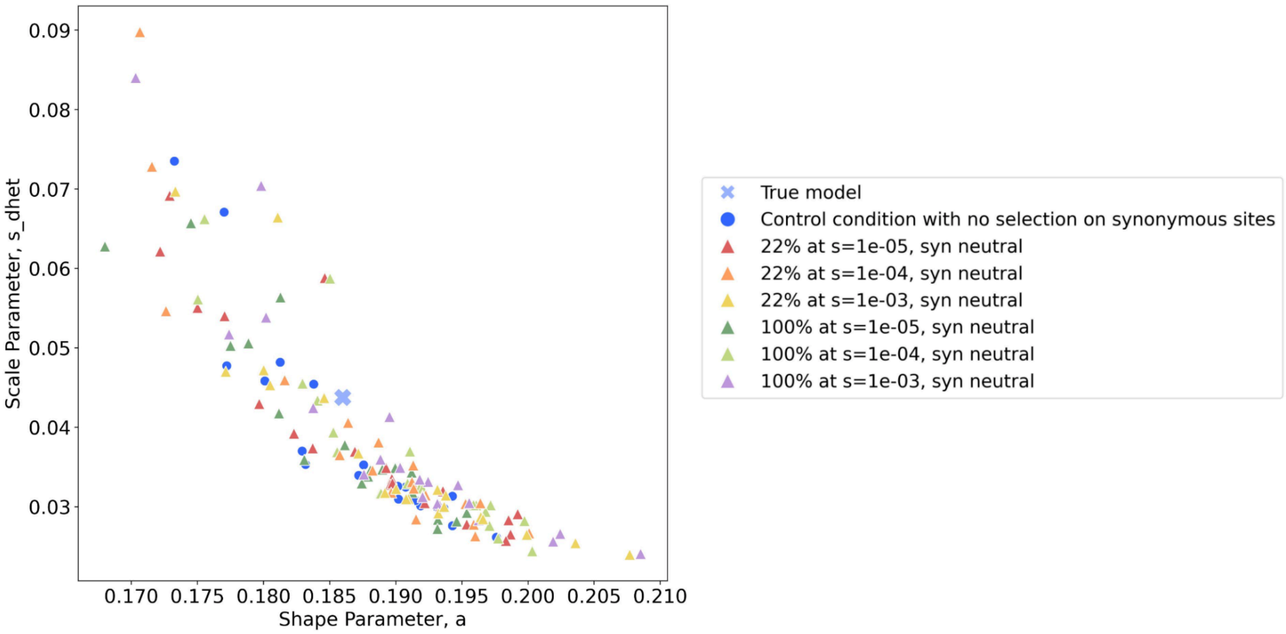


**Supplementary Figure 11: Distribution of inferred shape and scale parameters in a gamma DFE model for non-synonymous mutations when demography inferred from a known set of neutral variants.** Each point represents an individual simulation replicate. Scale parameter, $s_{dhet}$, represents the scale parameter in units of heterozygous selection strength.
